# Interpreting transient interactions of intrinsically disordered proteins

**DOI:** 10.1101/2022.11.14.516525

**Authors:** Samuel Wohl, Wenwei Zheng

**Affiliations:** Department of Physics, Arizona State University, Tempe, AZ 85287, USA; College of Integrative Sciences and Arts, Arizona State University, Mesa, AZ 85212, USA

## Abstract

The flexible nature of intrinsically disordered proteins (IDPs) gives rise to a conformational ensemble with a diverse set of conformations. The simplest way to describe this ensemble is through a homopolymer model without any specific interactions. However, there has been growing evidence that the conformational properties of IDPs and their relevant functions can be affected by transient interactions between specific and non-local pairs of amino acids. Interpreting these interactions from experimental methods, each of which is most sensitive to a different distance regime referred to as probing length, remains a challenging and unsolved problem. Here, we first show that transient interactions can be realized between short fragments of charged amino acids by generating conformational ensembles using model disordered peptides and coarse-grained simulations. Using these ensembles, we investigate how sensitive different types of experimental measurements are to the presence of transient interactions. We find methods with shorter probing lengths to be more appropriate to detect these transient interactions, but one experimental method is not sufficient due to the existence of other weak interactions typically seen in IDPs. Finally, we develop an adjusted polymer model with an additional short-distance peak which can robustly reproduce the distance distribution function from two experimental measurements with complementary short and long probing lengths. This new model can suggest whether a homopolymer model is insufficient for describing a specific IDP, and meet the challenge of quantitatively identifying specific transient interactions from a background of nonspecific weak interactions.

---

Intrinsically disordered proteins (IDPs) are notable for their ability to perform a variety of biological functions without taking on a well-defined structure.^1,2^ They are instead characterized by an ensemble of highly flexible conformations.^3–8^ Even in complex with other proteins, some IDPs can remain partially unfolded and dynamic, providing a balance between specificity and flexibility while maintaining high affinity.^9,10^ Recently, attention has been drawn to dynamic and disordered regions that exists in and out of the binding complex due to their potential role in regulating/assisting binding. Multiple studies show these flanking regions close to the binding site can alter the conformation and affinity of the complex.^11–13^ Possible explanations include direct interactions between flanking and binding regions or indirect interactions within an IDP that alter its conformational properties.^14,15^ In addition, some studies have suggested metastable interactions between specific amino acids far away from each other along an IDP sequence,^16–18^ which might significantly alter the conformational preference of an IDP from a random coil and regulate their biological function.^19,20^ Even under strong denaturing conditions, specific amino acid interactions were known to exist in folded proteins.^21^ There is therefore a strong motivation for investigating and quantifying these non-local, transient interactions within an IDP. This process faces substantial challenges both experimentally and computationally due to transient and flexible nature of these interactions.

The rapid growth of the IDP field has led to the development of many experimental techniques that can provide us with structural information. Some of these methods can offer indirect information on transient interactions by measuring local structural information, such as secondary structure information using circular dichroism (CD)^22^ or secondary structure chemical shift,^23^ and residual specific solvent accessibility using hydroxyl radical footprinting,^24^ ^19^F nuclear magnetic resonance (NMR)^25^ or solvent paramagnetic relaxation enhancement (sPRE).^26^ These methods require more in-depth analysis in combination with computational methods to provide details on transient interactions. Some other methods can directly measure distances between amino acids, including photoinduced electron transfer (PET),^27^ paramagnetic relaxation enhancement (PRE),^28^ Förster Resonance Energy Transfer (FRET),^29^ and small-angle X-ray scattering (SAXS),^30^ which are the main focus of this work. Each of these techniques is most sensitive to a specific distance regime, from here on referred to as the probing length. FRET measurements, for example, will only be able to accurately capture details close in distance to the chemical dye’s Förster radius,^31^ which ranges from ~4 to 7 nm. PRE and PET signals are more sensitive to a much shorter probing length than FRET, in the range from ~1.5 to 2.5 nm for PRE^32^ and from ~3 to 5 *Å* for PET.^27^

An IDP’s conformational ensemble usually has a very high degree of freedom due to its flexible nature. It is therefore not feasible to probe the entire ensemble using only experimental measurements. This has encouraged the use of computational methods that are capable of integrating multiple experimental techniques and molecular simulations to build a rigorous structural ensemble. One type of computational method is based on re-weighting the conformations of an initial ensemble in order to match experimental measurements.^4–7^ If the initial ensemble is reasonable, for example by using an optimal all-atom force field with sufficient sampling,^33^ ensemble fitting presents a comprehensive and cost-effective way for detailing the various conformations available to a given IDP. However, the method often has difficulty converging due to imperfect all-atom force fields,^34–36^ insufficient sampling,^37–39^ limited experimental measurements,^40^ and likely all of the above in its current state. A recent work has shown that the fitted ensemble relies heavily on the prior (initial ensemble).^41^ If the correct conformation has never been sampled in the initial ensemble, additional methods must be used, for example, to adapt conformations from initial ensemble to match the experimental measurements.^42^ Another method relies on directly biasing molecular simulations towards experimental measurements.^43,44^ This has proven to be very useful to introduce conformational changes in folded proteins,^45,46^ but is often cumbersome for IDPs. Experimental measurements of IDPs are ensemble-averaged properties and cannot be easily introduced as a biasing force in the simulation, often resulting in time consuming iterative procedures or maximum entropy principles.^47–50^ In addition, the outcome of the biased simulation is still dependent on the accuracy of the intial physics based model (i.e. force field) in the same way as ensemble fitting methods. Overall, the use of ensemble fitting and biased simulations to obtain IDP ensembles is still challenging at the moment. Improvements in all-atom force fields and growing sources of experimental data are expected to alleviate some of these concerns.

In the current stage where experimental measurements are limited, polymer models can be useful for identifying transient interactions since they have very few free parameters. Our recent efforts have shown that a self-avoiding walk model SAW-*ν*, in which a scaling exponent (*ν*) dependent distance distribution function *P*(*r, ν*) is used to adapt the variation of the chain dimension, can be used to analyze FRET and SAXS measurements and reconstruct an IDP’s distance distribution function.^51,52^ However, such a model only has a single peak in the distance distribution function, making it impossible to recreate transient interactions. A recent work from Pappu’s lab has shown that competing charge interactions lead to a bimodal end-to-end distance distribution and the distance distribution function can be modeled by an SAW model with an additional Gaussian function in the short-range regime.^53^ Motivated by this work, we plan to develop an improved SAW-*ν* model using this additional term to capture and resolve transient interactions from experimental measurements.

As shown in Fig. 1, in order to obtain sufficient data for parameterizing the new polymer model, we first design a series of model peptides with a few oppositely charged amino acids on each side(Fig. 1A). Coarse-grained simulations of these peptides suggest transient interactions exist between the two ends of the chain (Fig. 1B). These simulation data can help us narrow down free parameters in the distance distribution function. To further test if the new polymer model can reconstruct this *P*(*r*) using experimental measurements, we use these simulation data again to calculate artificial experimental measurements (Fig. 1C) and then obtain the model-interpreted *P*(*r*) by minimizing the difference between the artificial and model calculated experimental measurements (Fig. 1D). At last we can compare the model-interpreted *P*(*r*) (Fig. 1D) and the actual *P*(*r*) (Fig. 1B) using a statistical test to quantitatively assess our model. We would like to ask 1) how sensitive each type of experimental measurement are to the existence of transient interactions; 2) how many experimental measurements are needed to detect transient interactions; and 3) if an improved polymer model can be used to interpret transient interactions from artificial experimental measurements and reproduce the bimodal distance distributions.

**Figure 1:**
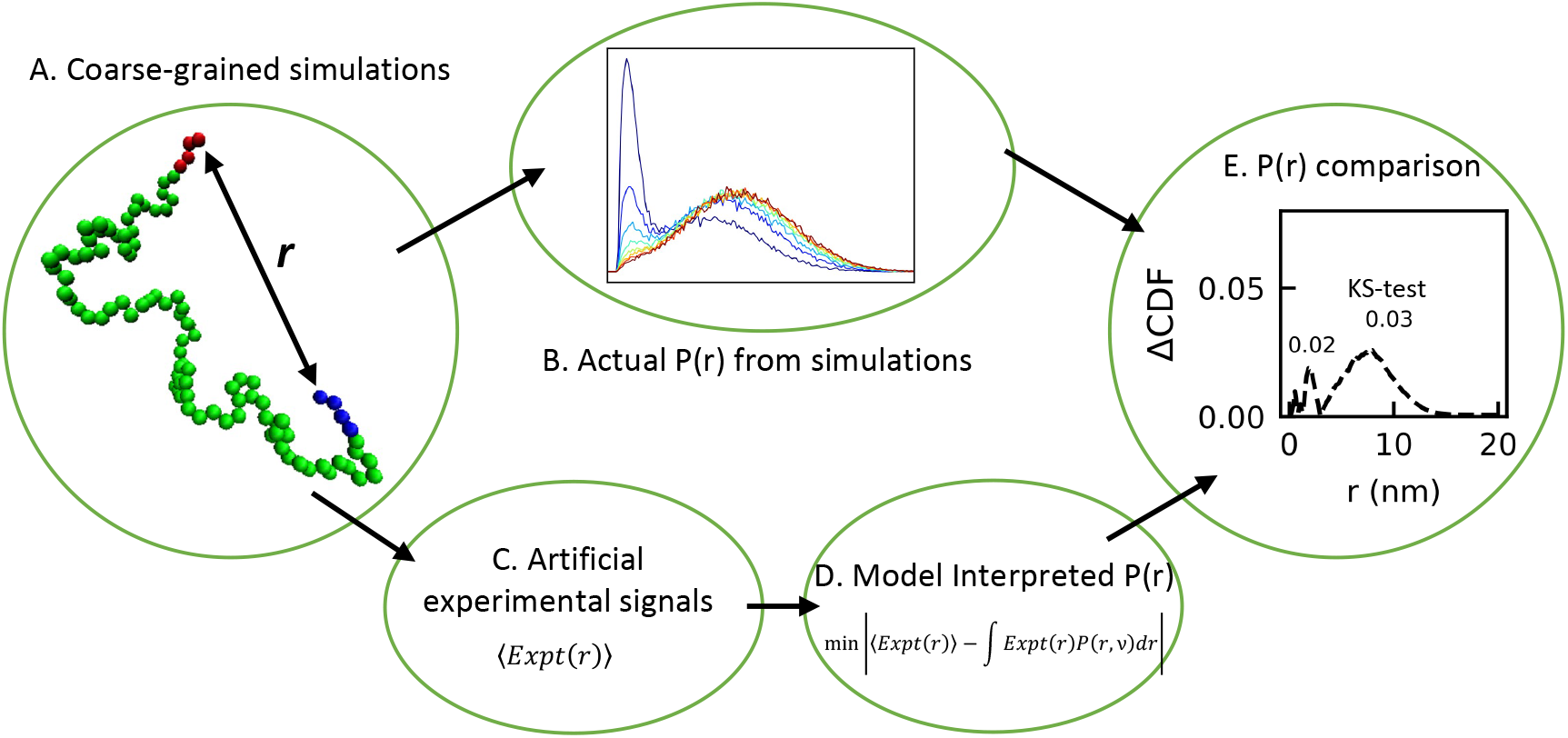
Schematic diagram of this work. A) Coarse-grained simulations of a model peptide with a few oppositely charged amino acids on the two sides. B) Distance distribution functions *P*(*r*) using end-end distances *r* obtained from coarse-grained simulations. C) Ensemble-averaging artificial experimental measurements calculated using the coarse-grained simulations. D) Minimizing the difference between the artificial experimental measurements and model calculated experimental measurements to obtain the model-interpreted *P*(*r*). E) Comparing the model interpreted and actual *P*(*r*) using the Kolmogorov-Smirnov (KS) test.

## Transient electrostatic interactions inside IDPs

Our first goal is to design a series of model IDPs which exhibit tunable transient interactions. It has already been shown that, for a charged sequence, different charge patterning can lead to bimodal behavior in the distance distribution function.^53^ However, for a typical IDP the fraction of charged amino acids are usually below 30%.^54^ It is not known what length of charged patch is necessary to introduce transient interactions nor how often these charged patches appear in an IDP. We therefore consider typical IDP sequences from DisProt database.^54^ We collect 530 sequences that were used in our previous work on the salting-out effect in IDPs,^55^ following two selection criteria: 1) only the longest disordered region is selected for each sequence and 2) only sequences between 30 to 400 amino acids were chosen to make them compatible with polymer theory and practical in a coarse-grained model. The frequencies of positively and negatively charged patches up to a length of eight residues are shown in Fig. 2(A). There is a clear downward trend between the probability of observing these charged patches and the length of the patches, *N*. Charged patches with more than eight residues are rare and negligible in these sequences. To further investigate the nature of these charged patches, we also calculate a reference line representing the probability to observe a charge patch with a length *N* assuming the amino acids are in random distributions but with the same amino acid composition (dashed lines in Fig. 2A). It is interesting that past *N* = 3, the probability of observing a patch starts to deviate from the reference line, rising slightly above it. The longer the charge patch, the larger the deviation. This suggests the naturally occurring IDP sequences are designed by evolution to have charge segregation, which can potentially lead to transient interactions in the conformational ensemble.

**Figure 2:**
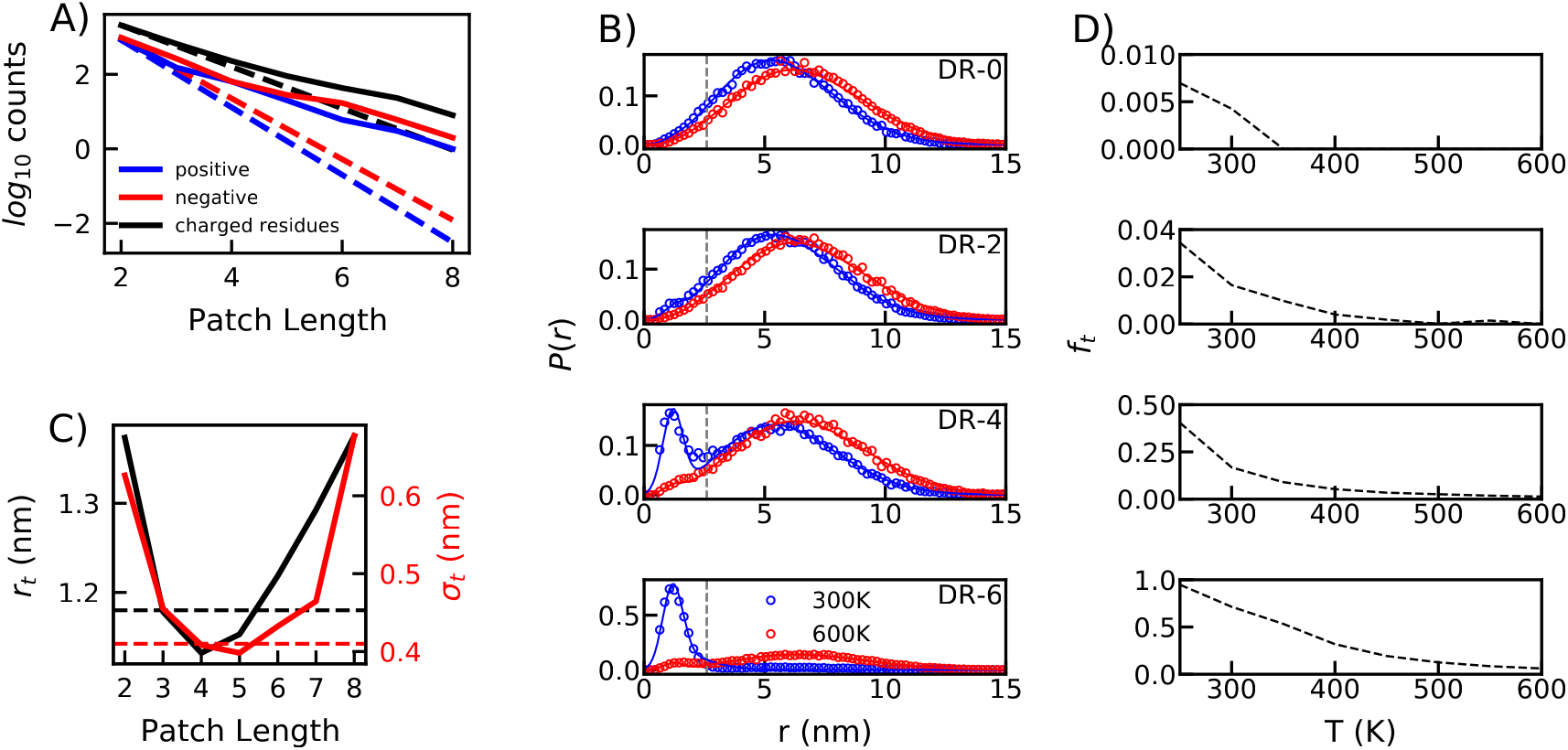
A) Prevalence of charged patches in IDPs from the DisProt database.^54^ Dashed lines indicate the expected counts if the amino acids in these sequences are randomly distributed. B) Simulated end-to-end distance distribution *P*(*r*) of DR-N at different temperatures *T* shown in the legends and patch lengths *N*. Solid lines indicate the best fit for each case using an adjusted SAW-*ν* model with an additional Gaussian distribution at the short-distance regime (Eq 1). Dash lines at *r*=2.6 nm show the separation of transient and polymer-like peaks. C) Fitting parameters and of the Gaussian distribution function (i.e. the center *r_t_* and the standard deviation *σ_t_*) to characterize transient interactions at different patch lengths. The values for a global fit for peptides from DR-2 to DR-6 are shown in dashed lines. D) The fraction of transient interactions from fitting Eq 1.

From this data we develop a minimalist model sequence that uses segregated positively and negatively charged amino acids to introduce transient interactions. The model IDP has a typical chain length of 100 residues, making it a good candidate for coarse-grained modelling. The two ends feature two charged patches, one consisting of *N* positively charged arginine and the other of *N* negatively charged aspartic acid. Glycine has limited interactions with the other amino acids, so it is only used as a spacer and for balancing the fraction of charged and uncharged amino acids. The series modeled peptide will be referred to as DR-N. Coarsegrained molecular dynamics simulations were performed, using the previously established HPS model,^56^ for patch lengths (*N*) of up to 8 charged residues and temperatures ranging from 250K to 600K for each peptide (see Supporting Methods for simulation details). Fig. 2B displays the *P*(*r*) for representative values of *N* at 300K (blue) and 600K (red) (see Supporting Fig. S1 for *P*(*r*) at other conditions). Note that the high temperatures here are not physiologically relevant and only acts as a method for adjusting the propensity of transient interactions. A metastable state exists at short-distance region for sequences with an *N* value larger than three at 300K, where electrostatic interactions between these charged patches pull the two ends of the chain together.

In order to quantify the fraction of transient interactions from these conformational ensembles, we examine how well a modified polymer model can describe the bimodal *P*(*r*). There are two components of the adjusted *P*(*r*) function,

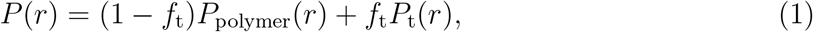

in which *f*_t_ determines the contribution of transient interaction term to the overall *P*(*r*). The first component *P*_polymer_(*r*) captures the polymer-like nonspecific weak interactions. Here we use our previously tested self-avoiding walk model with a varied scaling exponent *ν* (the SAW-*ν* model) to adjust the distribution function. The SAW-*ν* model has been shown to be able to effectively interpret both the polymer scaling exponent *ν* and reconstruct the distance distribution from FRET^51^ or SAXS experiments.^52^ The distribution function was written as

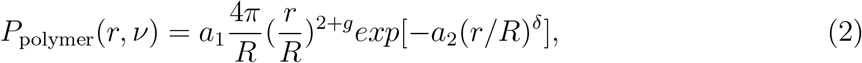

in which *g* = (*γ* – 1)/*ν*^57^ and *γ* =1.1615 in three dimensions,^58^ *δ* = 1/(1 – *ν*),^59^ and the constants *a*_1_ and *a*_2_ are normalization factors so that 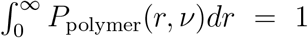 and 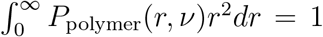. The second component *P*_t_(*r*) captures the short-distance transient interactions. We use a Gaussian distribution function similar to a recent work for capturing the bimodal behavior of the distance distribution function for charged peptides,^53^ which can be written as

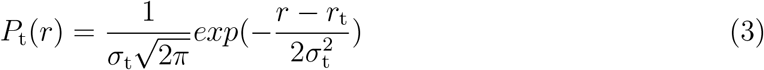

in which the center *r*_t_ and standard deviation *σ*_t_ are free parameters. Together with the scaling exponent *ν* and the weight of transient interaction term *f*_t_, the overall adjusted *P*(*r*) has four free parameters and can be used to fit the *P*(*r*) from conformational ensembles of DR-*N* peptides.

For each sequence of DR-*N* with conformational ensembles at a variety of temperatures *T*, we minimize the root means squared difference between the simulated *P*(*r*) and modeled *P*(*r*) using Eq. 3 to find the optimum values of *r*_t,*N*_,*σ*_t,N_, *f*_t,*N,T*_, and *ν_N,T_*. Note we assume transient interactions should be peptide (sequence) dependent, and therefore *r*_t,*N*_ and *σ*_t,*N*_ are set to be the same for the same peptide at all temperatures whereas *f*_t,*N,T*_ and *ν_N,T_* are different for the same peptide at different temperatures. Fig. 2C shows the range of *r*_t_ and *σ*_t_ values found by fitting the adjusted *P*(*r*) in Eq. 1 from DR-2 to DR-8 at different temperatures. *r*_t_ decreases from 1.37 nm at *N* = 2 to 1.13 nm at *N* = 4 and increases again as *N* increases. *σ*_t_ follows the same parabolic behavior but reaches its minimum at *N* = 5. The uncertainty of *r*_t_ and *σ*_t_ is large for *N* smaller than 2 due to the low fraction of transient interactions and are therefore not shown here. We suspect that it might be possible to predetermine *r*_t_ and *σ*_t_ and reduce the number of free parameters in the new model since neither values appear to depend strongly on patch length *N*. In order to obtain a general set of *r*_t_ and *σ*_t_, we employ a global fitting for all sequences from DR-2 to DR-6. DR-7 and DR-8 are not included because patches that size do not appear often in IDPs, as shown in Fig. 2A. To minimize the summation of the root mean squared (RMS) difference between the 40 *P*(*r*) distributions from five sequences each with eight ensembles at different temperatures and their corresponding modeled *P*(*r*), there are a total of 82 free parameters. The first 80 parameters are *ν* and for each case and the last two are the global values for *r*_t_ and *σ*_t_. Dashed lines in Fig. 2C show the global fitting results of *r*_t_ and *σ*_t_ as 1.18 and 0.41 nm, respectively. A fixed set of *r*_t_ and *σ*_t_ makes the adjusted polymer model flexible enough to describe *P*(*r*) across a range of ensembles with different fraction of transient interactions while reducing the number of free parameters. The fraction of transient interactions reduces as *N* decreases or temperature increases, and can be completely negligible for all sequences at 600K (Fig. 2D). At 300K, DR-4 has well balanced transient and polymer-like interactions and will therefore be used as a representative case for further investigation.

## Transient interactions from diverse experimental measurements

From the coarse-grained simulations of the DR-4 peptide, we obtain a series of conformational ensembles with different fractions of transient interactions ranging from ~40% at 300K to ~0% at 600K (see Fig. 2D and Supporting Fig. S1). This provides a good data set for testing how well a variety of experimental technologies can detect the existence of transient interactions from a background of weak polymer-like interactions. The advantage of using a coarse-grained simulation data set for such a test is two-fold. First, we can easily calculate the experimental signals directly from the molecular trajectories by averaging over a larger pool of diverse conformations. Second we also have the true reference physical properties such as the distance between a pair of amino acids, the polymer scaling exponent, and the radius of gyration relevant to different experimental technologies. The simplest way to perform such a test is then to 1) calculate the artificial experimental signal from simulations, 2) obtain the model-interpreted physical properties such as *ν* and/or *P*(*r*) using the old SAW-*ν* model, 3) and compare the model-interpreted properties with simulated properties (see Fig. 1). A larger deviation indicates that transient interactions have a greater impact on the homopolymer interpretation of the experimental method and the technique is more sensitive to, or equivalently more appropriate to detect, transient interactions.

To start, we use FRET as an example. In a FRET experiment, a donor and an acceptor dye are covalently linked to two specific amino acid sites of interest along the chain. The donor dye is optically excited and the excited energy can either be emitted as a photon or transferred to an acceptor dye. The energy transfer efficiency *E*_FRET_(*r*) is related to the distance *r* between the pair of dyes, if the dye can rapidly experience different orientations within time scales of donor life time. The first step in analyzing FRET’s sensitivity to transient interactions is to convert DR-4 simulation data to an artificial FRET efficiency. For each frame in the simulation, we consider the labels to be at two ends of the peptide and calculate the end-to-end distance r. In dotted symbols of Fig. 3A, we show the distance distribution *P*(*r*) obtained from the simulation at two representative temperatures 300K and 600K with different *f_t_*. At 600K, transient interactions are negligible and there is one single peak along *P*(*r*), commonly seen in a homopolymer model.^60–62^ We can then calculate the FRET efficiency for each frame of the trajectory using the Förster equation^31^ as

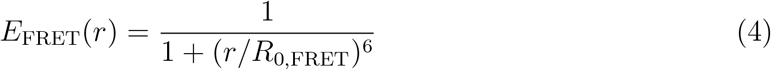

in which *R*_0,FRET_ is the Förster radius, a value intrinsic to a given set of dyes. This value determines the center of the sigmoid shape function between *E*_FRET_(*r*) and *r* (see the inset of Fig. 3A) and can be thought as the probing length of FRET, since it determines what distance regime the FRET measurement is most sensitive to. By varying the type of dyes, *R*_0,FRET_ can range from 4.5 to 6.5 nm. For testing the method, we use *R*_0,FRET_ as 5.4 nm, meaning values of *r* far above or below 5.4 nm do not impact the FRET efficiency. The ensemble averaging FRET efficiency can then be thought as an artificial FRET measurement to our simulated conformational ensemble.

**Figure 3:**
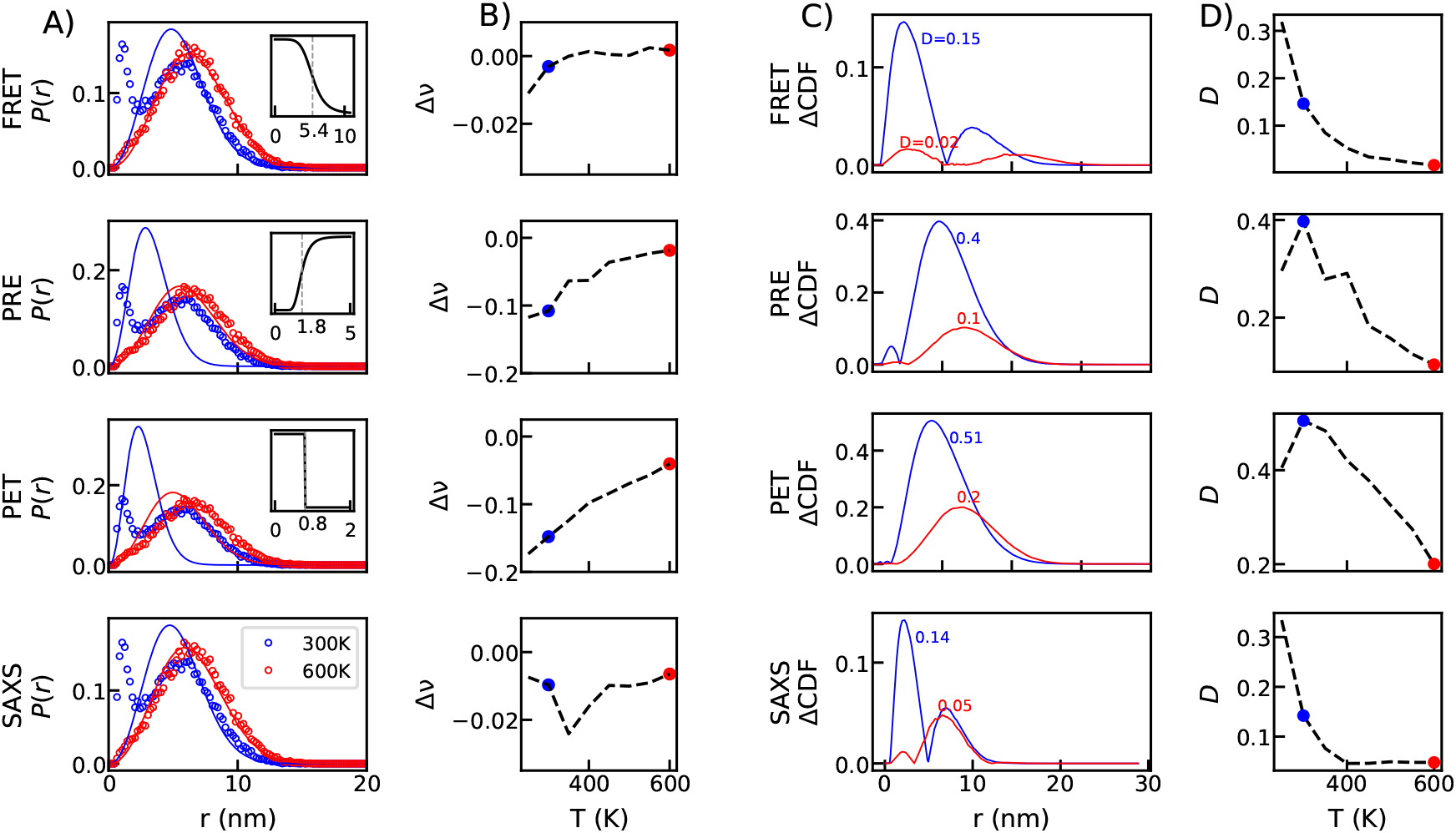
Using the SAW-*ν* model to interpret transient interactions from a variety of experimental signals. A) Comparison between the model-interpreted (solid lines) and simulated P(r) distributions (circles) from four different experimental methods for DR-4. Colors show the two representative temperatures with 300K in blue and 600K in red. B) Δν defined as the difference between the model-interpreted *ν* and actual *ν* as a function of simulation temperatures. C) Difference between the model-interpreted and simulated cumulative distribution functions, with the maximum *D* value (KS test) as labelled. D) *D* as a function of simulation temperatures.

Once the artificial FRET efficiency has been determined, we move on to the second step: reconstructing the structural properties using our recently developed SAW-*ν* model.^51^ In this model, there is only one free parameter, the polymer scaling exponent *ν*, in the distance distribution function *P*(*r, v*). By minimizing the difference between model interpreted FRET efficiency and artificial FRET efficiency, we find the *P*_model_(*r*) obtained using a polymer model (red lines in Fig. 3) is in excellent agreement with the simulated *P*(*r*) (red dotted symbols). This is consistent with our observation that transient interactions do not exist in the CG simulation at 600K. However, at 300K the *P*(*r*) is bimodal, similar to the distribution seen in a recent work using sequences with mostly charged amino acids.^53^ The single peak SAW-*ν* model cannot capture either of the two peaks in *P*(*r*), suggesting that the FRET interpreted model is affected by the presence of transient interactions.

We further develop two metrics to quantify the deviation between the structural properties as interpreted by SAW-*ν* model and from the simulated reference. This effectively tells us what level of transient interactions cause the old homopolymer model to break down, or similarly how well a given experimental method can detect transient interactions. The first metric Δ*ν* is the difference between the model-interpreted *ν*^51^ and the *ν* that can be directly calculated from the simulation trajectory by fitting the RMS distance as a function of the sequence separation.^63^ As shown in Fig. 3B, Δ*ν* is negative for most temperatures, indicating that transient interactions cause the SAW-*ν* model to underestimate the scaling exponent. The deviation becomes more prominent for simulated ensembles at lower temperatures when transient interactions become more prominent. Δ*ν* predictably increases towards zero as the temperature rises and transient interaction subsides.

Since *ν* is only well defined for a homopolymer, it becomes problematic to interpret *ν* in a conformational ensemble with transient interactions, which can be considered heteropolymeric behavior. We therefore develop a second metric by directly quantifying the deviation between the model-interpreted and simulated distance distribution functions. We have performed the Kolmogorov-Smirnov (KS) test on the cumulative distribution function (CDF) as shown in Fig. 3C, which should give a better indication of whether model-interpreted *P*(*r*) fails to capture transient interaction peak. For analyzing FRET, the maximum deviation *D* always occurs at the short-distance peak position. This is due to FRET’s probing length (~5.4 nm) being relatively large compared to the center of transient-interaction peak (~1.2 nm). Since the *P*(*r*) in the SAW-*ν* model has only one peak, FRET’s large probing length encourages the model to capture only the long-distance regime. The model interpreted *P*(*r*) is still slightly affected by the existence of transient interactions and shifts towards the short-distance peak. Similar to Δ*ν*, *D* approaches zero as transient interactions reduce with temperature. We see qualitatively the same behaviors between Δ*ν* and *D* when interpreting FRET measurement across conformational ensembles with different transient interactions (Fig. 3B and D).

The same framework can be applied to investigating how sensitive common experimental techniques are to transient interactions. We further investigated three other methods which can provide distance information, paramagnetic relaxation enhancement (PRE), photoinduced electron transfer (PET), and small-angle X-ray scattering (SAXS). In a PRE experiment, the measured intensities of the cross peaks for the paramagnetic and diamagnetic forms of the protein can be calculated as^32,64^

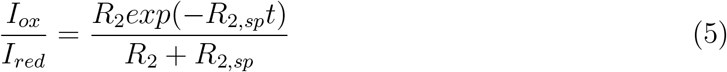

in which *t* is the total evolution time of the transverse proton magnetization and *R*_2_ is the intrinsic relaxation of the proton. Here we use *t*=10.87ms and *R*_2_ = 10/s for the test. *R*_2,*sp*_ is the contribution to the relaxation caused by the paramagnetic probe and therefore is relevant to the distance *r* between the paramagnetic probe and the backbone amide proton. The relation between *R*_2,*sp*_ and *r* can be described by

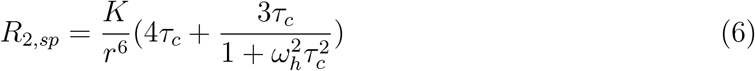

in which *K* is a property of the spin label, *ω_h_* is the Larmor frequency of the proton spin and *τ_C_* is the apparent PRE correlation time dependent on the molecular weight of the protein. Here we use *K* = 1.23 × 10^-32^*cm*^6^/*sec*^2^ for MTSL label,^65^ *ω_h_*=850 Mhz and *τ_C_* = 5 ns for the test. As shown in the inset of Fig. 3, the relation between the PRE signal (*I_ox_/I_red_*) and the distance *r* is also in a sigmoid shape similar to FRET. There is no direct analog to the FRET probing length *R*_0,FRET_ for PRE, but we find 1.8 nm to be a good estimate for our test set of parameters for PRE signal. In principle both *R*^2^ and *τC* are sequence dependent and the probing length of PRE can approximately range from 1.5 to 2.5 nm, still much smaller than the range of *R*_0,FRET_ for using different FRET dyes. PRE’s short probing length makes the method more sensitive to the presence of transient interactions than FRET. As shown in Fig. 3B, Δ*ν* still approaches zero as temperature increases, but is overall with a much larger deviation than for FRET measurements. This suggests that, since the probing length of PRE is more in line with the short-distance transient interaction, the PRE interpretation of the old polymer model is greatly hampered by transient interactions. Much like Δ*ν*, *D* values from PRE observations are larger than those from FRET at every temperature. Interestingly, *D* peaks at 300K and then decreases with temperature, as opposed to the monotonic decrease previously observed. Unlike FRET, interpreting PRE with the old model can capture the *P*(*r*) at the lowest temperature where transient-interaction peak is dominant (see Supporting Fig. S2). The highest deviations are then seen when a conformational ensemble has equally prominent transient and polymer-like peaks at 300K.

In a PET experiment, the relation between the PET rate and the distance *r* can be described by a step function *k*_0_*H*(*r* – *R*_0,PET_) in which *k*_0_ is the maximum quenching rate.^27,66^ *R*_0,PET_ can be considered as the probing length, the distance regime at which PET is expected to be most sensitive. For testing the method using a coarse-grained simulation with residue-level resolution, we set *R*_0,PET_ to be 0.8 nm. The value of *k*_0_ will not affect the test since it will be canceled out when interpreting the artificial PET rate. As shown in Fig. 3B and *D*, Δ*ν* and *D* as measured from PET follow roughly the same trends as those from PRE. At 300K, the single peak from the model-interpreted *P*(*r*) is in the middle of transient-interaction and polymer-like peaks of the simulated *P*(*r*), leading to a large deviation. The deviation eventually approaches zero as transient interactions decrease. Once again, *D* peaks at 300K due to the equally prominent transient-interaction and polymer-like peaks. Notably, Δ*ν* and *D* are larger for PET based observations than those from PRE, despite their relatively similar probing lengths. This is probably due to the fact that *R_p_* for PET lies below transient-interaction peak position, making it even more sensitive to such interactions.

The last method we would like to test is SAXS. Even though SAXS does not directly provide the distance between a specific pair of amino acids, the SAXS intensity does depend on the pairwise distance distribution between every pair of atoms in the molecule. It would be interesting to check whether it is more close to FRET with a large probing length less sensitive to transient interactions or PRE and PET with a small probing length more sensitive to these interactions. The overall framework has to be modified slightly since SAXS does not directly depend on the distance between a pair of amino acids. We first calculate the SAXS intensity from the coarse-grained conformation using a set of form factors *f_aa_* developed for 20 amino acids^67^ as

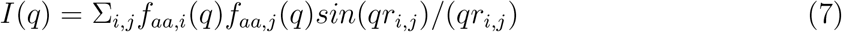

in which *r_i,j_* is the distance between *i*-th and *j*-th amino acids and q is the scattering vector. The ensemble averaging *I*(*q*) is then obtained for each conformational ensemble at different temperatures (see Supporting Fig. S3). In order to interpret these artificial SAXS intensities we use Guinier analysis to obtain the radius of gyration *R_g_*.^68^ The scaling exponent *ν* can be obtained using the SAW-*ν* model providing the relation between *R_g_* and *ν*.^51^ With this model-interpreted *ν*, we can create the model-interpreted *P*(*r, ν*). Δ*ν* from SAXS across different ensembles has a unique behavior in contrast to the other three experimental methods, remaining mostly small from 250K to 600K with a maximum deviation at 350K. We attribute this behavior to the fact that SAXS is most sensitive to the overall size of the protein instead of a well defined probing length. If the *P*(*r*) is close to a single peak at lowest and highest temperatures, model-interpreted *ν* is reasonable. The largest variation happens when *P*(*r*) has two peaks and *ν* does not correlate with the size of the protein any more. However what is more interesting is that *D* does not look the same as Δ*ν* but follows the same trend as FRET. This is mainly due to the fact that at the lowest temperature with only transient interaction peak in the *P*(*r*) (see Fig. S1), the SAW-*ν* model cannot capture this distance distribution but *ν* interpreted using the same model is still reasonable to capture the size of the protein.

In order to explore general behavior between the probing length of an experimental technique and its ability to detect transient interactions, we consider a generic experiment with a sigmoid shape signal as a function of amino acid distance similar to FRET (Eq. 4), but with an adjustable probing length, *R_p_*, substituted for *R*_0,FRET_. Following the same analysis framework and using the SAW-*ν* model, we can then obtain Δ*ν* and *D* values for a range of *R_p_* as a function of temperature, or equivalently the fraction of transient interactions, as shown in Fig. 4. At high temperatures, when transient interactions become negligible, both Δ*ν* and *D* are close to zero for any *R_p_* larger than 2 nm. This suggests any experiment with a reasonably large *R_p_* can capture polymer-like interactions centered at approximately 6 nm (Fig. S1). At lower temperatures and for higher fractions of transient interactions, the model tolerates a smaller range of *R_p_*. Δ*ν* is minimized only when *R_p_* is ~6 nm, suggesting the scaling exponent is more heavily influenced by the polymer-like peak than the transient-interaction one. *D* behaves differently at the lowest temperature since the two metrics capture different physical properties. Δ*ν* is based on the polymer scaling exponent, making it most sensitive to the polymer-like, weak interactions. *D* captures the difference in *P*(*r*) instead of *ν*, and therefore has no preference for the polymer-like or transient-interaction peak. At the lowest temperature, with about 40% transient interactions, Δ*ν* remains small when *R_p_* is close to the center of the polymer-like peak while *D* accurately reflects the difference between the model-interpreted and simulated *P*(*r*) at the transient-interaction peak. Since *ν* can be ill-defined within a conformational ensemble that deviates from a homopolymer, we believe *D* is a better metric for checking how well the adjusted model captures transient interactions.

**Figure 4:**
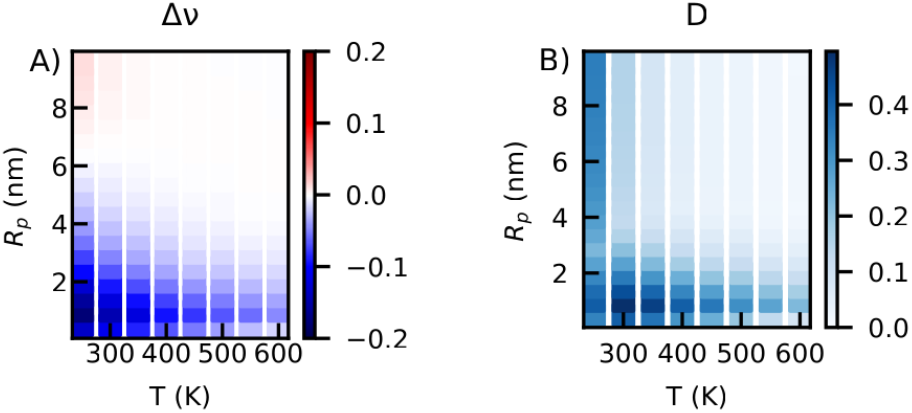
Δ*ν* (A) and *D* values (B) as a function of the probing length *R_p_* and the temperature *T* quantifying fraction of transient interactions inside the DR-4 ensembles.

With this new probing length-based view of experimental methods, we can break down the previously tested experimental methods into two groups. Both FRET and SAXS are methods with relatively large probing lengths and are very effective at capturing the overall size of an IDP and its polymer-like interactions. They are both useful for obtaining *ν*, as shown previously,^51,52,69^ and can reasonably estimate *ν* even in the presence of transient interactions, as shown in Fig. 4B. However, this makes them poor predictors of transient interactions. Meanwhile, PRE and PET can be considered methods with relatively small probing lengths and exhibit the opposite behaviors. Increasing transient interactions disrupts their ability to characterize physical properties like *ν* or the overall size of an IDP, while at the same time making them ideal for detecting these non-local behaviors. What we would like to further investigate is whether an adjusted polymer model can reconstruct the *P*(*r*) distribution with both transient-interaction and polymer-like peaks when we combine short- and long-range experimental techniques.

## Interpreting transient interactions using an adjusted polymer model

As shown in the previous section, the current SAW-*ν* model with its single-peak *P*(*r*) cannot capture both transient and polymer-like interactions at the same time. The adjusted polymer model shown in Eq. 1, including an additional short-distance peak, would be more appropriate, but introduces additional free parameters and necessitates the use of more than one experimental restraint. In principle, there are four free parameters in the *P*(*r*) of Eq. 1: *ν*, which determines the shape of the polymer-like interaction peak (Eq. 2); and f_t_, *r*_t_, and *σ*_t_ which determine the fraction, location and width of the transient-interaction peak, respectively (Eq. 3). The optimization is then underdetermined using only one or two experimental restraints. We therefore propose to investigate whether an adjusted polymer model with fixed values of *r*_t_ and *σ*_t_ (i.e. 1.18 and 0.41 nm), obtained from globally fitting all DR-2 to DR-6 ensembles as discussed in the previous section, can interpret transient interactions. This model will be referred to as SAW-*ν*-tr.

The first question we would like to ask is whether the SAW-*ν*-tr model can reconstruct *P*(*r*) by a combination of two experimental technologies. Using a combination of FRET and PET as an example, we can follow the same scheme as shown in Fig. 1. We first calculate the artificial FRET and PET signals for every ensemble of DR-4 at different temperatures, second obtain the two fitting parameters *ν* and *f*_t_ by minimizing the RMS difference between the SAW-*ν*-tr model-interpreted and simulated *P*(*r*), and finally quantify the difference between the model-interpreted and simulated *P*(*r*) using the *D* value from KS test. We need to adjust the KS test to identify which of the two peaks along *P*(*r*) are being properly fit since both transient-interaction and polymer-like peaks need to be tested. We know from existing simulations that *P*(*r*) has a local minimum at approximately 2.6 nm as shown in Fig. 2B, and therefore choose to identify the maximum difference between the two CDFs above and below that point with two additional metrics, *D*_1_ and *D*_2_. *D*_1_ is the maximum difference between the two CDFs when *r*¡2.6 nm and indicates how well the reconstructed *P*(*r*) matches transient-interaction peak. Similarly, *D*_2_ comes from *r*≥2.6 nm and indicates how well the polymer-like peak is reproduced. Using the new SAW-*ν*-tr model to match measurements of FRET and PET improves the reconstruction of *P*(*r*), with *D*_1_=0.06 and *D*_2_=0.04 (the first row of Fig. 5). In contrast, the single-peak SAW-*ν* model returned *D*=0.15 for FRET and *D*=0.51 for PET as shown in Fig. 3. However, FRET and PET together still overestimate the presence of transient interactions and underestimate polymer-like interactions. This is probably due to the rather small probing length of PET, making the method sensitive to even shorter distance regime than transient interactions seen in the DR-4 simulations.

**Figure 5:**
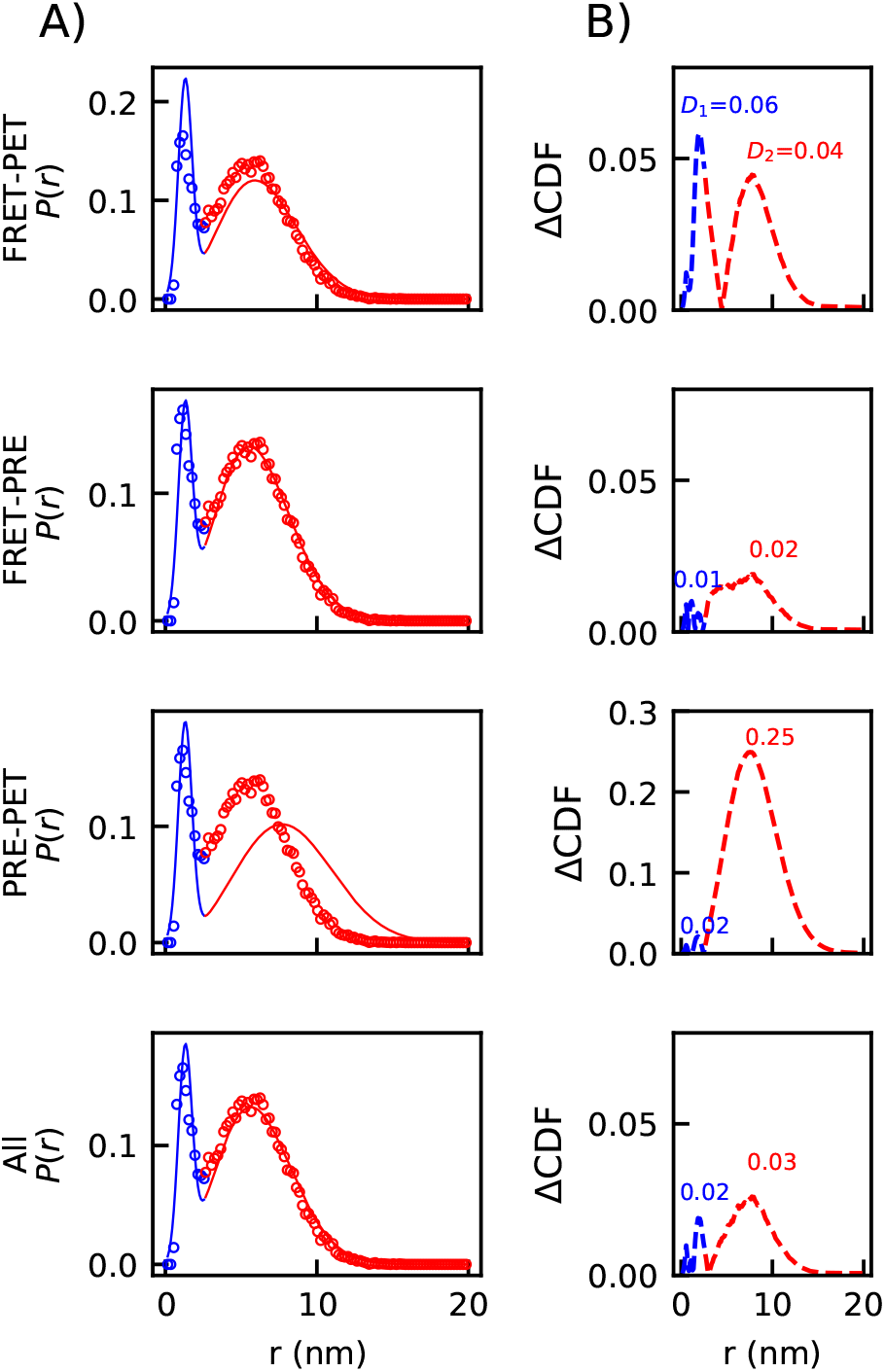
Using the SAW-*ν*-tr model to interpret transient interactions from a combination of different experimental methods. A) Comparison between the model-interpreted (solid lines) and simulated *P*(*r*) distributions (circles) from four different combinations of experimental methods for DR-4 at 300K. Transient-interaction peak is shown in blue and the polymer-like peak is shown in red. B) Difference between the model-interpreted and simulated cumulative distribution functions quantified using the maximum difference within *r*¡2.6 nm (blue, *D*_1_) and *r*≥2.6 nm (red, *D*_2_).

The combination of FRET and PRE (the second row of Fig. 5), on the other hand, makes for a more accurate interpretation than the combination of FRET and PET. In addition to reconstructing both peaks with very small *D*_1_ = 0.01 and *D*_2_ = 0.02, this combination also successfully captures the minimum position of *P*(*r*) at 2.6 nm. It therefore benefits from the slight larger probing length of PRE, compared to PET, which is closer to the center of transient-interaction peak. The PRE and PET combination (the third row of Fig. 5) leads to a low *D*_1_ value of 0.02 but a much higher *D*_2_ value at 0.25. Omitting the long-range experimental technique corresponding to the center of the polymer-like peak makes the SAW-*ν*-tr model unable to reproduce the long-distance regime of the *P*(*r*). Finally, we consider whether the prediction can be further improved by including all three experimental measurements at once (the fourth row of Fig. 5). By combining FRET, PRE, and PET, the reconstruction actually becomes slightly worse, with both *D*_1_ and *D*_2_ increased by 0.01, in contrast to only using a combination of FRET and PRE. This is mainly due to the fact that there are only two peaks in the simulated *P*(*r*) and therefore to capture both these peaks, two experimental restraints are sufficient. For real applications of a specific sequence, additional experimental restraints might suggest the addition of a third peak within *P*(*r*). However, this requires empirical knowledge regarding the position and width of the third peak.

Finally we would like to understand what combination of experimental methods in general can be used with the SAW-*ν*-tr model to reproduce the *P*(*r*) including both transient and polymer-like interactions. Similar to the previous case where we investigated one experimental input(Fig. 4), we employ two generic experimental signals. Both signals have a sigmoid shape as a function of *r*, similar to FRET (Eq. 4), but have two independent probing lengths, *R*_*p*,1_ and *R*_*p*,2_, substituted for the corresponding *R*_0,*FRET*_. We apply the SAW-*ν*-tr model to reconstruct the *P*(*r*) of the DR-4 ensemble at 300K with balanced transient and polymer-like interactions using these two experimental inputs. The test to the other temperatures with different fraction of transient interactions are shown in Supporting Fig. S4. *D*_1_ and *D*_2_ are still calculated for each case to measure how well transient-interaction and polymer-like peaks have been reproduced, respectively, but we also consider the overall reconstruction as described by *D*: the larger of *D*_1_ and *D*_2_. As shown in Fig. 6, *D*_1_ is minimized when at least one experiment with a probing length close to transient-interaction peak (i.e. approximately 2 nm) is used, regardless of the second *R_p_* value. *D*_2_ is minimized when one *R_p_* is within the range of 3 to 5 nm and the other is within the range of 4 to 8 nm, both close to the polymer-like peak (i.e. approximately 6 nm). The diagonal however is always extremely large, suggesting one experimental method, even analyzed by the newly developed SAW-*ν*-tr model, cannot capture both transient-interaction and polymer-like peaks in the *P*(*r*). This is due to the fact that, in order to robustly optimize the two free parameters in the SAW-*ν*-tr model, one needs at least the same number of experimental restraints. Together with the previous analysis using one experimental input with the old SAW-*ν* model (Fig. 4), we can therefore conclude that one experimental method is not sufficient to quantify the fraction of transient interactions due to the existence of polymer-like interactions.

**Figure 6:**
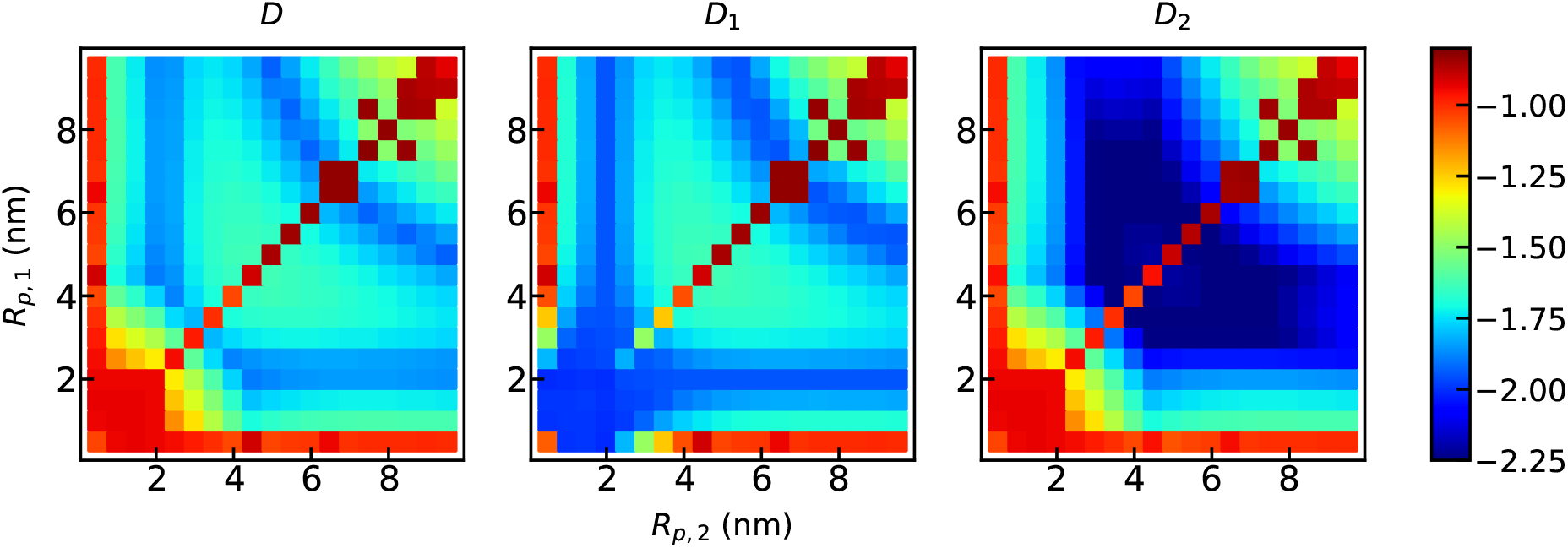
*D* (A), *D*_1_ (B) and *D*_2_ (C) as a function of the probing lengths *R*_*p*,1_ and *R*_*p*,2_ from two experimental technologies when trying to reconstruct the *P*(*r*) using the SAW-*ν*-tr model. Color bar is in log_10_ scale.

When looking at the overall reconstruction quality using *D*, we see the first minimum with one *R_p_* between 1.5 and 2.5 nm and the other *R_p_* larger than 4 nm. This agrees with our expectation and previous observation when combining FRET and PRE that two complementary experimental technologies, with one small and one large probing length corresponding to the transient interaction peak and polymer-like peak respectively, form the best combination to characterize the overall *P*(*r*). Interestingly, there also exists a second minimum for pairs of probing lengths both sitting close to the polymer-like peak. In this case, the fitting is expected to properly match the size of the polymer-like peak, but can also subsequently reweight transient interaction peak as well. Since in the SAW-*ν*-tr model the balance between the two peaks is controlled by one free parameter, adding one additional experimental restraint with a probing length at a different regime of the polymer-like peak is sufficient to determine both *ν* (shape of the polymer-like peak) and the fraction of the polymer-like peak together. This interesting observation expands the range of experimental technologies that are able to determine both transient and polymer-like interactions. For instance, two FRET measurements at the same pair labeling positions, but using different pairing dyes with different Förster radii, are in principle able to reconstruct the double-peak *P*(*r*) with the help of the SAW-*ν*-tr model.

At last we would like to investigate the robustness of scheme when varying the *r*_t_ and *σ*_t_ values. It is not possible to simultaneously determine these values at the same time using the SAW-*ν*-tr model without more experimental restraints. We test a variety of *r*_t_ and *σ*_t_ and perform the same reconstruction test over a range of probing lengths. As shown in Supporting Fig. S5, when *σ*_t_ is fixed at 0.4 nm and *r*_t_ changes from 1.0 to 1.4 nm, *D*_2_ is not affected since the adjustment is only made to the Gaussian function used to capturing transient-interaction peak. *D*_1_ is lowest when *r*_t_=1.2 nm, close to the optimal value we determined. It is still reasonably small for *r*_t_=1.1 and 1.3 nm. Similar results are shown when *σ*_t_ is changed to 0.35 (Fig. S6) and 0.45 nm (Fig. S7). These tests suggest that the method with a fixed set of *r*_t_ within range of 1.1 to 1.3 nm and *σ*_t_ within range from 0.35 to 0.45 nm can still provide a reasonable estimate for *P*(*r*) and the fraction of transient interactions. In practice, transient interactions might not be caused by solely charged amino acids and therefore the location and width of transient interaction peak can differ from what the coarse-grained model proposes. We therefore further apply the same test using different ensembles of DR-2, DR-3, DR-6 and DR-8 (see Supporting Fig. S8). We show that even for these peptides in which the intrinsic *r*_t_ and *σ*_t_ are quite different (Fig. 2C), the scheme still works reasonably well, suggesting the method remains robust even when the location and width of the transient-interaction peak in the model might not be exactly the same as the reality.

## Conclusion

Homopolymer models such as Gaussian chain and self-avoiding walk models were commonly used for understanding the conformational properties of IDPs. These methods rely on the assumption that specific interactions are limited or negligible and nonspecific weak interactions dominate within an IDP. However, there is growing experimental evidence that there can be transient interactions between specific pairs or groups of amino acids within an IDP. In this work, we showed that, in a coarse-grained simulation of a sequence with two short fragments of charged amino acids, transient interactions exist. The distance distribution function therefore cannot be described by a homopolymer model such as the SAW-*ν* model with a single peak in the distribution function. A survey of typical IDP sequences suggested such short charged patches are not uncommon.

We therefore investigated what experimental methods can detect these interactions and how many are necessary. We found that experiments with smaller probing lengths, the characteristic distance at which the experimental signal changes the most, are more affected by the existence of transient interactions in an IDP conformational ensemble. However, one experiment cannot resolve both transient interactions and polymer-like weak interactions at the same time, and an additional experimental method with a different probing length can help. Given the complexity of integrating two experimental technologies, we introduced an adjusted polymer model, SAW-*ν*-tr model, in which one additional Gaussian distribution function has been added to the original SAW-*ν* model. We found that the new model is able to successfully reconstruct the distance distribution function containing both transient-interaction peak in the short-distance regime and polymer-like peak in the long-distance regime, by integrating two experimental technologies with complementary probing lengths. The entire framework therefore provides a simple yet effective scheme to quantitatively detect the heteropolymeric behavior of an IDP from existing experimental measurements and further identify specific transient interactions from a background of nonspecific weak interactions.

It remains to be seen in what manner transient interactions will appear in more complicated sequences. Functional IDPs may feature shorter and more frequent charged patches featuring a variety of sequence separations. Hydrophobic amino acids and especially aromatic ones might also be able to introduce transient interactions. All these aspects can complicate the current analysis and are worth further investigation.

## Acknowledgement

This work is supported by the National Science Foundation (MCB-2015030) and the National Institutes of Health (R35GM146814). The authors acknowledge Research Computing at Arizona State University.

