## Supporting Information for "Interpreting transient interactions of intrinsically disordered proteins"

### Supporting Information for: "Inferring transient interactions of intrinsically disordered proteins"

**Coarse-grained simulations**

The coarse-grained HPS model[1] considers each amino acid as one bead, described by three types of interactions: bonded, electrostatic and short-range pairwise interaction terms characterized by amino acid hydropathy.[2] Bonded interactions are modeled by a harmonic potential with a spring constant of 10 kJ/Å$^2$ and a bond length of 3.8 Å. Electrostatic interactions are modeled using a Coulombic term with Debye-Hückel [3] electrostatic screening,

$$E_{ij}(r) = \frac{q_i q_j}{4\pi D r}\exp(-r/\kappa), \tag{S1}$$

in which $\kappa$ is the Debye screening length and $D = 80$, the dielectric constant of the solvent. The short-range pairwise interaction is modeled using Ashbaugh-Hatch functional form[4],

$$\Phi(r) = \begin{cases} \Phi_{LJ} + (1-\lambda)\epsilon, & \text{if } r \leq 2^{1/6}\sigma \\ \lambda\Phi_{LJ}, & \text{otherwise} \end{cases} \tag{S2}$$

in which $\Phi_{LJ}$ is the standard Lennard-Jones potential

$$\Phi_{LJ} = 4\epsilon\Big[\Big(\frac{\sigma}{r}\Big)^{12} - \Big(\frac{\sigma}{r}\Big)^{6}\Big]. \tag{S3}$$

The $\lambda$ value in the pairwise interaction term is the arithmetic average of the $\lambda$ values of the two corresponding amino acids. The amino-acid specific parameters of the model are shown in Table S1. The interaction strength $\epsilon$ is set to 0.2 kcal/mol based on parameterization from previous works.[1]

Table S1. The amino acid parameters used in the model. $\sigma$ is the diameter of the amino acid used in the short-ranged pair potential. $\lambda$ is the scaled hydropathy from the literature [2].

| Type | Mass (amu) | Charge | $\sigma$ (Å) | $\lambda$ |
|---|---|---|---|---|
| ARG | 156.20 | 1 | 6.56 | 0.000 |
| ASP | 115.10 | -1 | 5.58 | 0.378 |
| GLY | 57.05 | 0 | 4.50 | 0.649 |

The first 1000 frames of each simulation are disregarded since they included remnants of the initial nonphysical conformation. That leaves 49000 frames each with an end-to-end distance $r$. $P(r)$, the distance distribution, is a histogram of the $r$ values from a given simulation separated into a bin width of 0.1 nm.

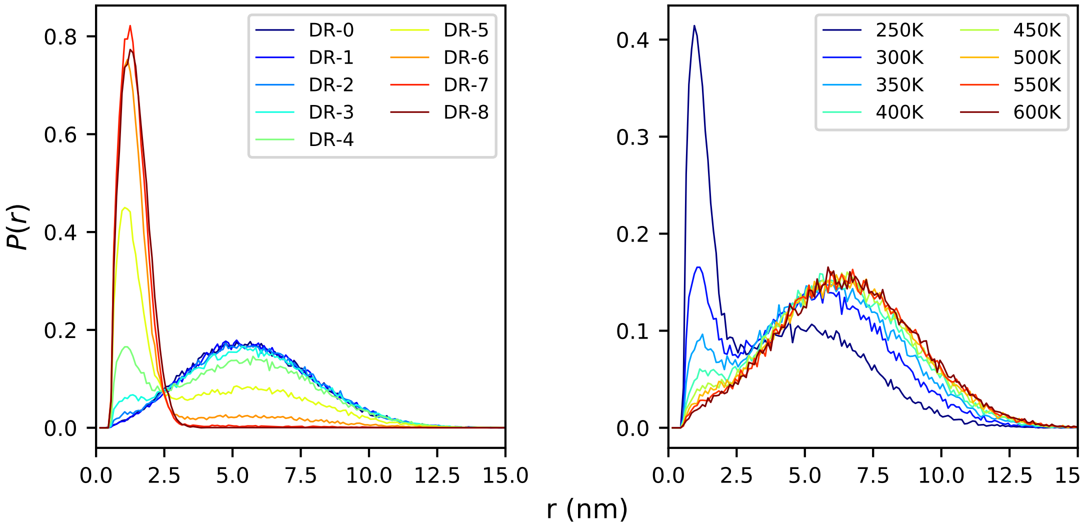


Figure S1. End-to-end distance distribution for sequences DR-0 through DR-8 at 300K (left) and for DR-4 at temperatures 250K through 600K (right).

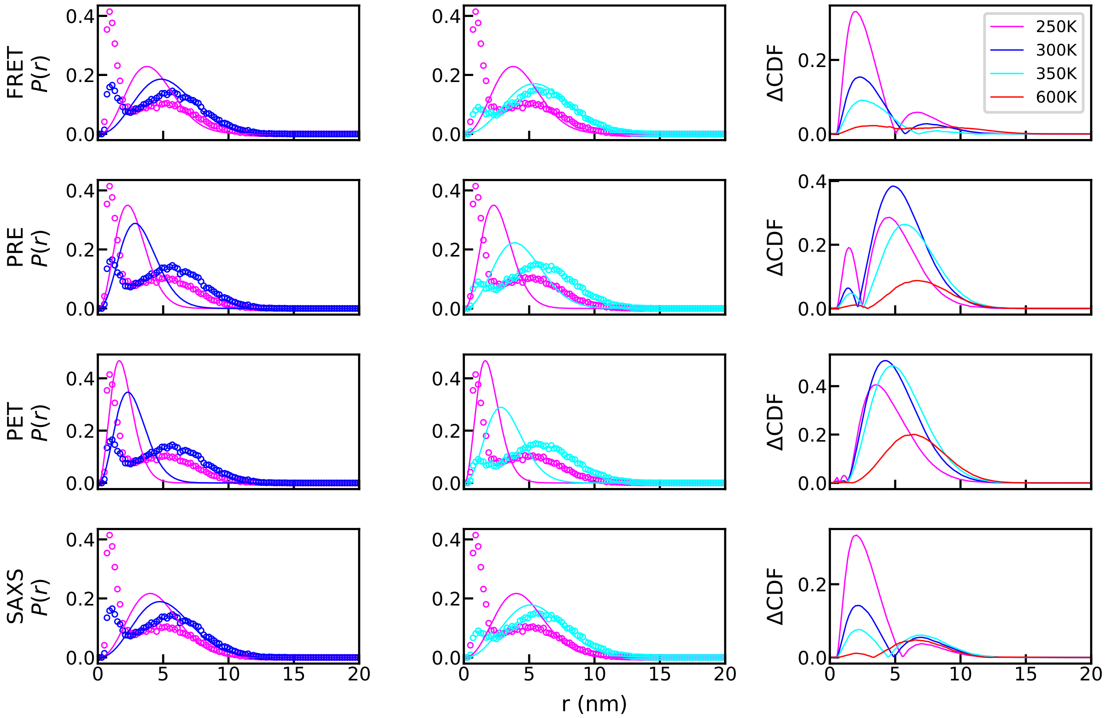


FIGURE S2. Using the SAW-$\nu$ model to interpret the transient interactions from a variety of experimental signals. Additional temperatures of interest (as shown in legends) are included to illuminate the difference between model-interpreted and simulated $p(r)$.

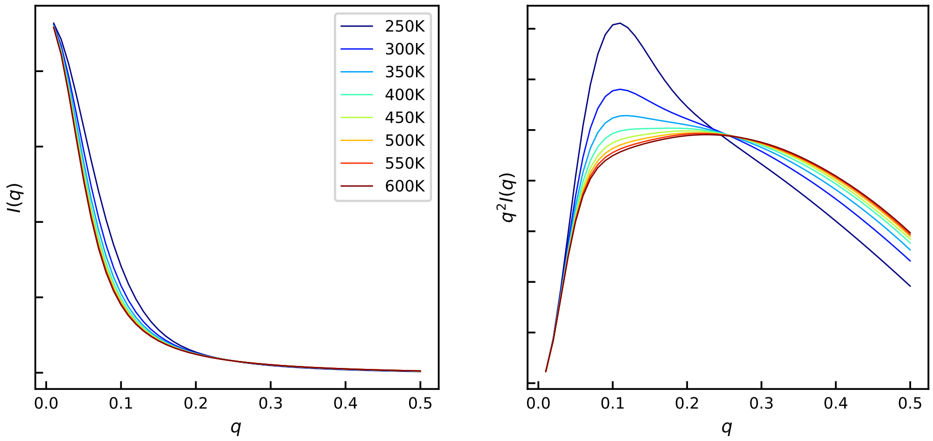


FIGURE S3. SAXS profile (left) calculated using the DR-4 trajectories at a range of temperatures as well as the corresponding Kratky plot (right).

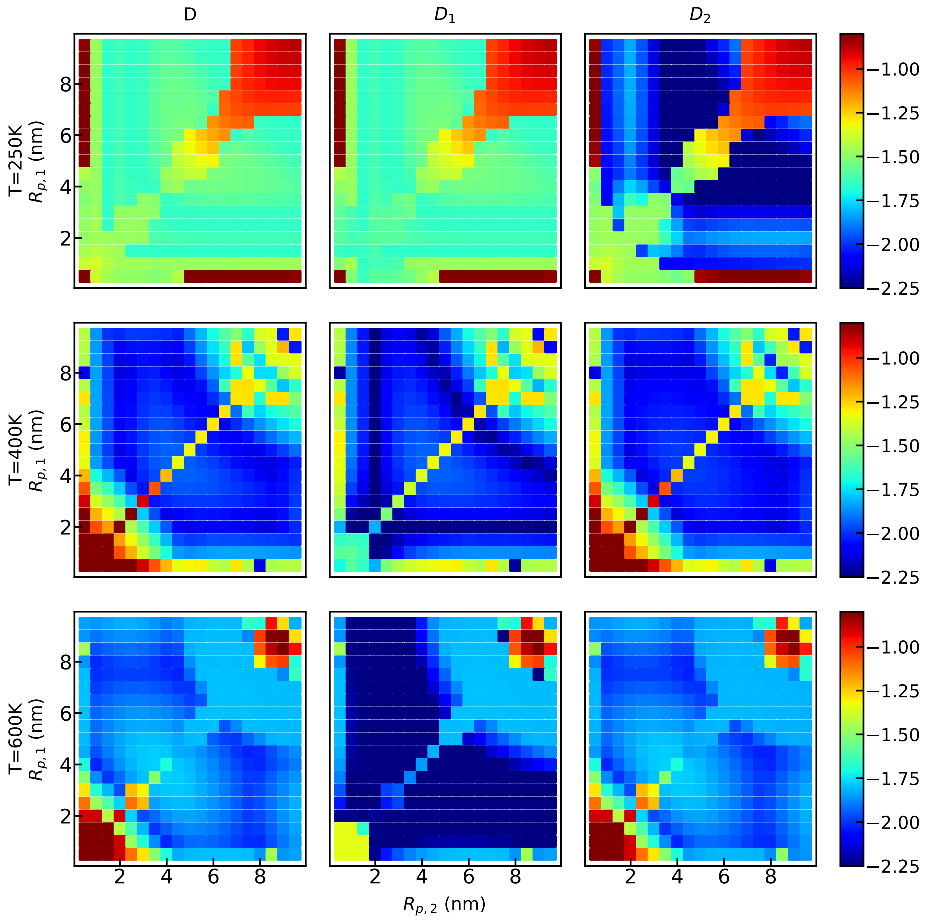


FIGURE S4. $D$, $D_1$ and $D_2$ as a function of the probing lengths $R_{p,1}$ and $R_{p,2}$ from two experimental technologies when trying to reconstruct the $P(r)$ using the SAW-$\nu$-tr model. DR-4 ensembles at multiple temperatures are tested. Color bar is in $log_{10}$ scale.

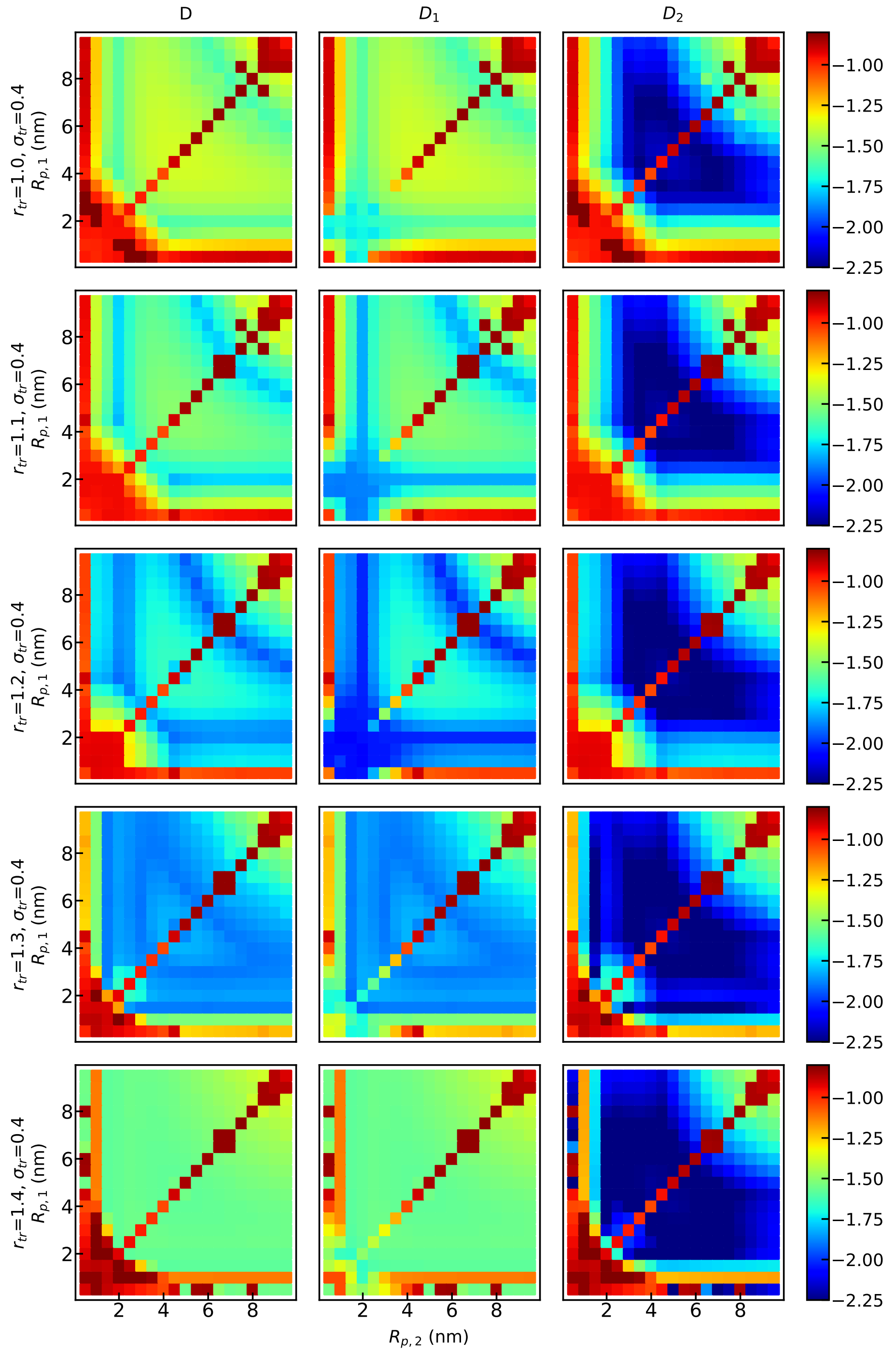


FIGURE S5. $D$ (left), $D_1$ (middle) and $D_2$ (right) as a function of the probing lengths $R_{p,1}$ and $R_{p,2}$ from two experimental technologies when trying to reconstruct the $p(r)$ using the SAW-$\nu$-tr model. $\sigma_{\rm tr}$ is set to be 0.4 nm and $r_{\rm tr}$ is scanned from 1.0 to 1.4 nm. Color bar is in $log_{10}$ scale.

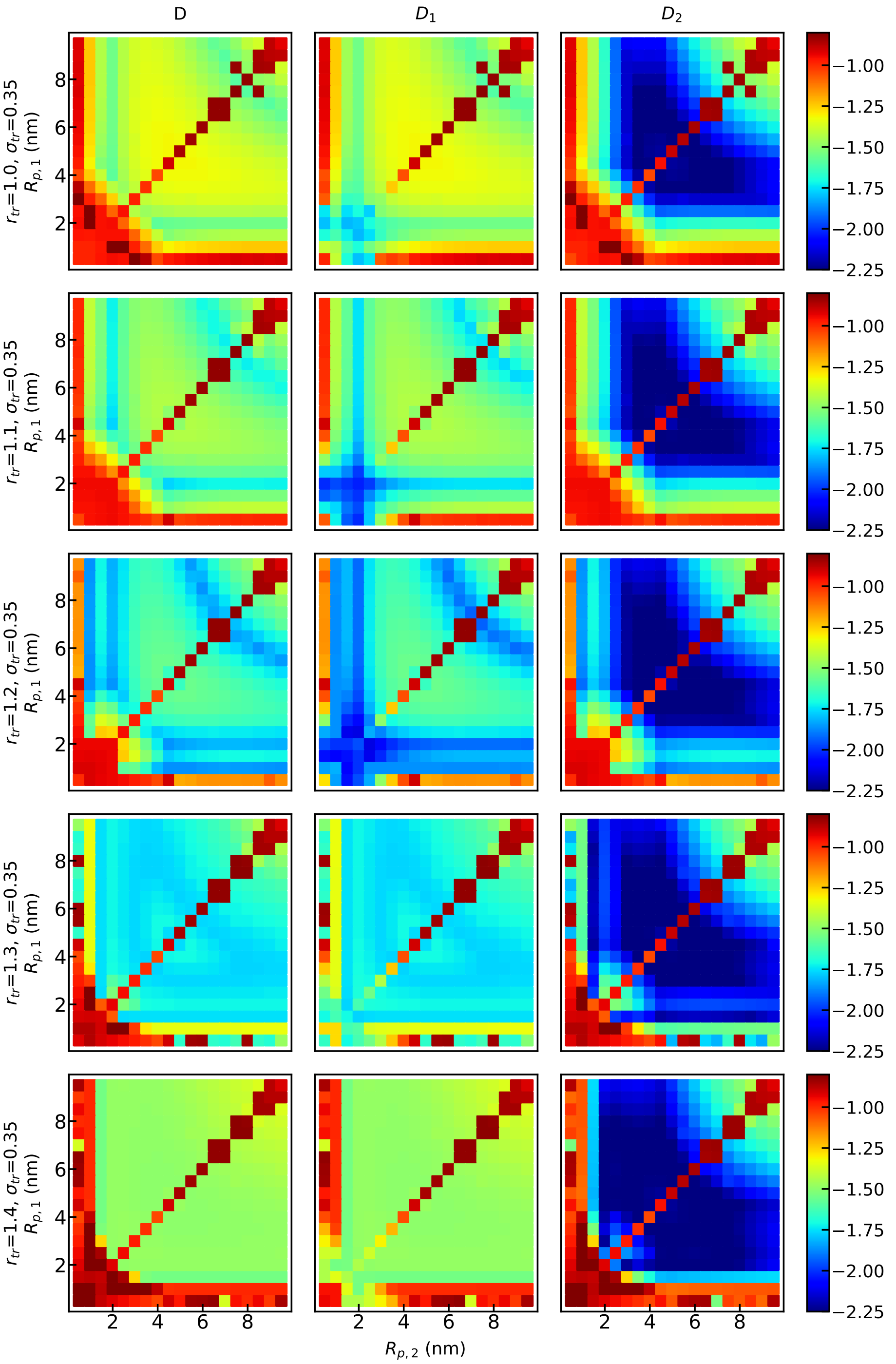


FIGURE S6. $D$ (left), $D_1$ (middle) and $D_2$ (right) as a function of the probing lengths $R_{p,1}$ and $R_{p,2}$ from two experimental technologies when trying to reconstruct the $p(r)$ using the SAW-$\nu$-tr model. $\sigma_{\text{tr}}$ is set to be 0.35 nm and $r_{\text{tr}}$ is scanned from 1.0 to 1.4 nm. Color bar is in $log_{10}$ scale.

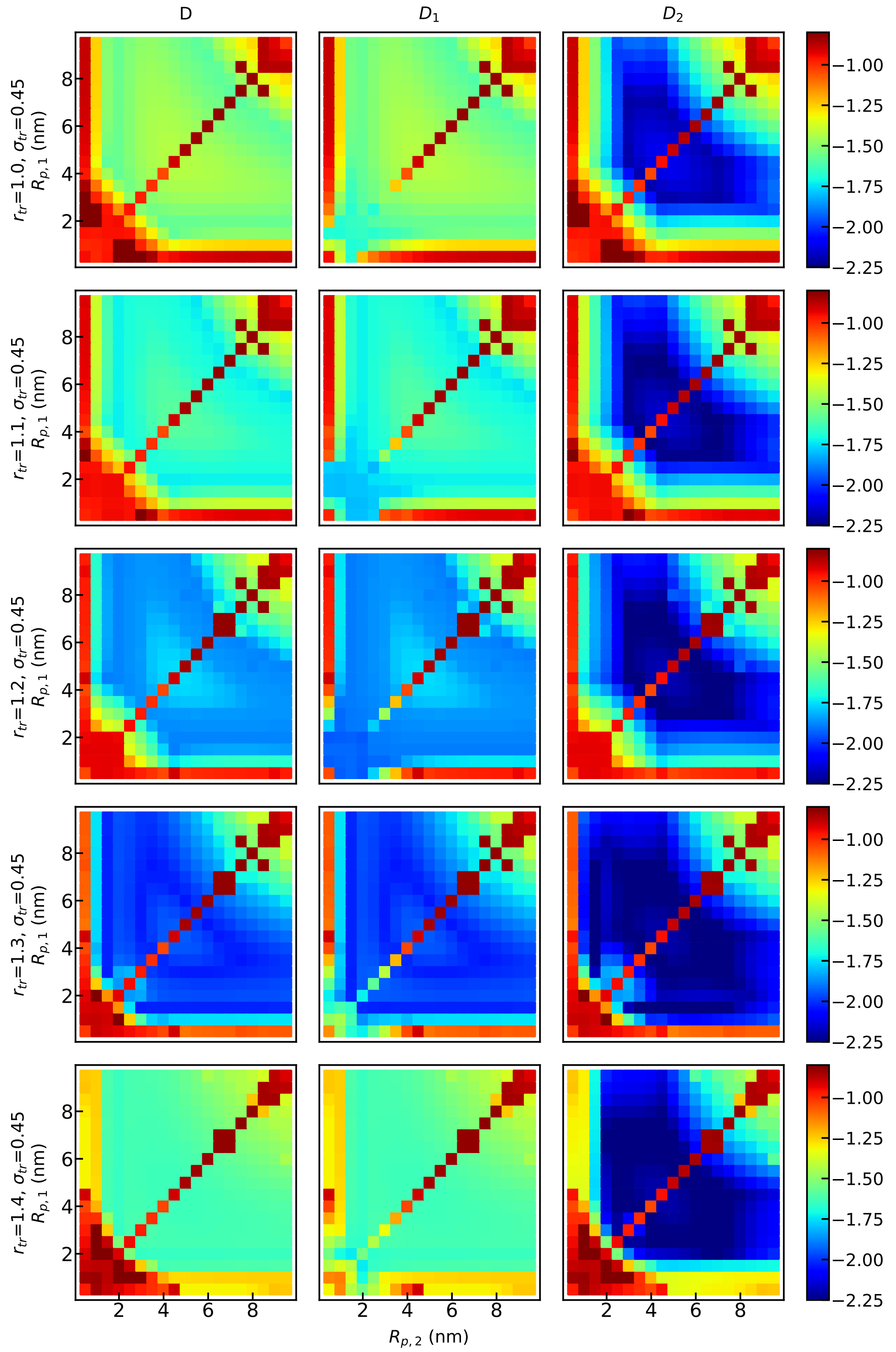


FIGURE S7. $D$ (left), $D_1$ (middle) and $D_2$ (right) as a function of the probing lengths $R_{p,1}$ and $R_{p,2}$ from two experimental technologies when trying to reconstruct the $p(r)$ using the SAW-$\nu$-tr model. $\sigma_{\text{tr}}$ is set to be 0.45 nm and $r_{\text{tr}}$ is scanned from 1.0 to 1.4 nm. Color bar is in $log_{10}$ scale.

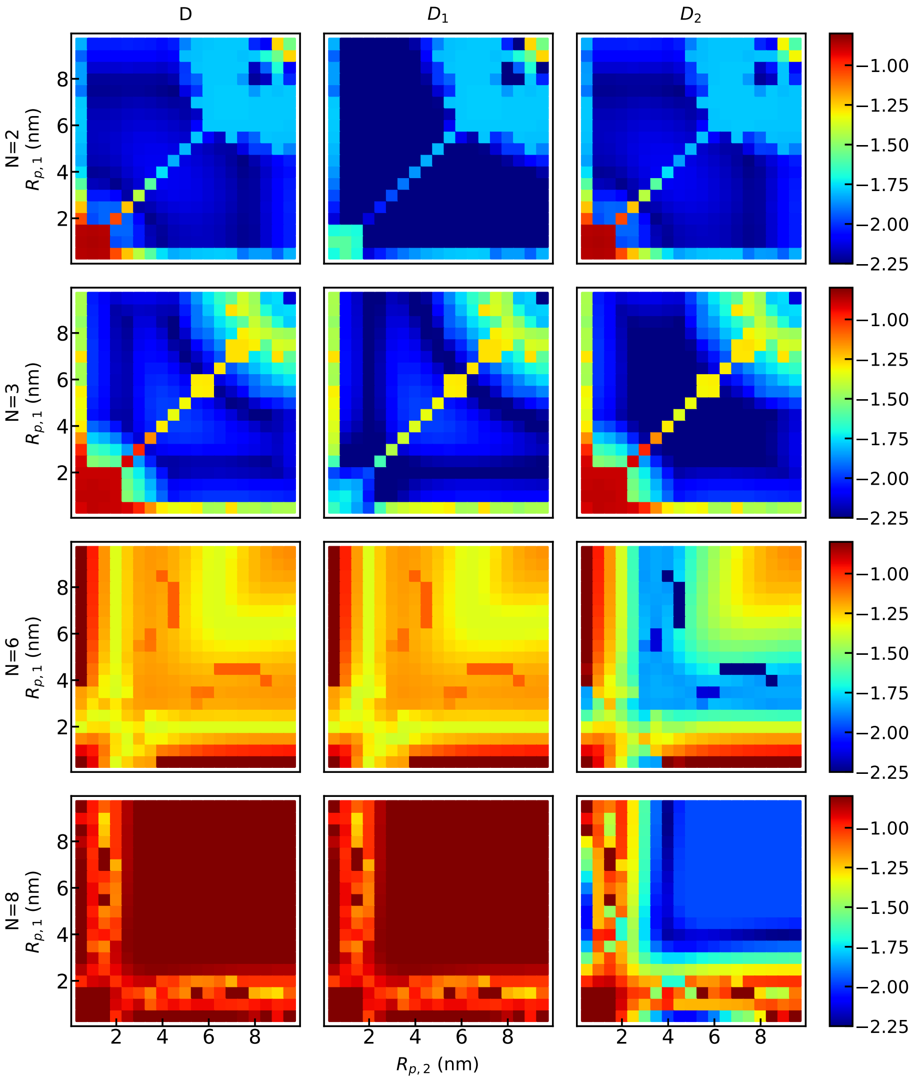


Figure S8. $D$ (left), $D_1$ (middle) and $D_2$ (right) as a function of the probing lengths $R_{p,1}$ and $R_{p,2}$ from two experimental technologies when trying to reconstruct the $P(r)$ using the SAW-$\nu$-tr model. DR-2, DR-3, DR-6 and DR-8 ensembles at 300K are tested. Color bar is in $log_{10}$ scale.
